## Supporting Information for "Cytoplasmic flow induced by a rotating wire in living cells: Magnetic rotational spectroscopy and finite element simulations"

**Outline**

**S1** – Experimental setup: Magnetic rotating field device and associated equipment

**S2** – Example of a field of view of a HeLa cells monolayer incubated with magnetic wires

**S3** – Sampling wires according to their length

**S4** – Review of the literature data on the intracellular viscosity obtained for MCF-10A, MCF-7 and MDA-MB-231 breast epithelial cells

**S5** – Comparison between the Newton and Oldroyd-B models based on finite element simulations

**S6** – Relative variation of the velocity component along $Oy$ and of the velocity modulus in the $Ox$, $Oy$, and $Oz$ directions

**S7** – COMSOL Multiphysics finite element simulations of rotating wires in the conditions $L/D >>1$

**S8** – Evidence of power-law relationships in MRS data across the studied cell lines for $D(L)$, $L^{*}(L)$, $\omega_{C}(L^{*})$ and $\eta_{0}\left( \omega_{C} \right)$

Complementary information including:

- Movies (.avi) illustrating key steps in data acquisition, such as wire entry into living cells, wire rotation within the cytoplasm
- COMSOL simulations of wire rotation in Newton and Oldroyd-B viscoelastic fluids
- COMSOL simulation files (.mph) for selected wire rotation cases, along with representative raw velocity field data (.txt)
- Cytoplasmic viscosity and elasticity data (for the four cell lines analyzed are also included and accessible through the same link
- Wire calibration data using water-glycerol mixture of known viscosity

is available via the ZENODO repository, with the link: <https://doi.org/10.5281/zenodo.17335446>.

**Keywords:**

Cell microrheology | Intracellular viscosity | Finite element simulations | Viscoelasticity | Magnetic Rotational Spectroscopy

This version Tuesday, December 16, 25

Revised version submitted to Journal of the Royal Society Interface

**Supporting Information S1**

Experimental setup: Magnetic rotating field device and associated equipment

The rotating magnetic field used for wire actuation is generated by a custom-built device consisting of four electromagnetic coils, each composed of 745 turns of copper wire (diameter 0.5 mm) wound in 17 layers [1]. The coils are arranged orthogonally in pairs at 90° to each other around the sample chamber. Their cores are made of 80:20 nickel-iron mu-metal, a soft ferromagnetic material with high magnetic permeability, characterized by a saturation magnetization of 0.7 tesla and a low coercivity below 2 A m⁻¹. Each aligned coil pair can generate magnetic fields ranging from 0 to 30 mT at the sample position. The electrical signals powering the coils (resistance 4.68 ohms) are supplied by a low-frequency function generator (arbitrary waveform, 50 MHz, two channels) and a homemade current amplifier. A rotating field is obtained by applying a phase shift of 90° between the signals sent to each orthogonal pair of coils. To control the temperature of the cell sample during experiments, a stream of air is directed toward the chamber through an inlet integrated into a thermally insulated cover. The system allows thermalization of the sample environment between room temperature and 50 °C [1].

Microscopy is performed using an inverted optical microscope equipped with ×20, ×40, ×60, and ×100 objectives, enabling both bright-field and phase contrast imaging (**Fig. S1**). The numerical apertures of the objectives are 0.4, 0.55, 0.7, and 1.3, respectively. Image acquisition is carried out using a QImaging EXi Blue camera controlled by Metamorph software. The coils and thermal cover are mounted on a supporting structure made from aluminum and poly(vinyl chloride) components for mechanical stability and ease of alignment.

Wire suspensions are processed prior to use with a 40 kHz, 110 W sonicator bath, which is employed to reduce wire lengths to a few micrometers when required for specific intracellular measurements. Samples are mounted using Gene Frame dual adhesive spacers to form sealed observation chambers. For data analysis, time-lapse image sequences are processed using ImageJ software and associated plugins, which enable manual or automated tracking of wire orientation over time.

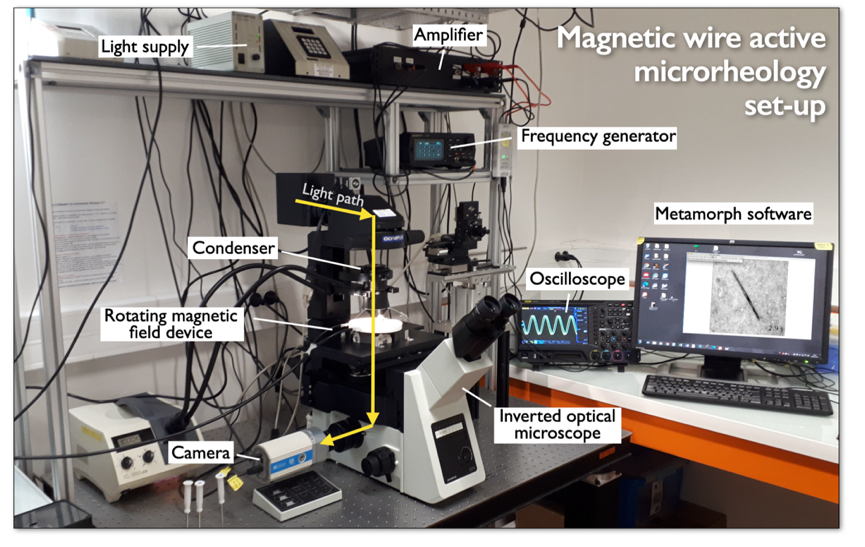

**Figure S1:** Optical microscopy set-up used to perform magnetic rotational spectroscopy (MRS) experiments.

**Supporting Information S2**

Example of a field of view of a HeLa cells monolayer incubated with magnetic wires

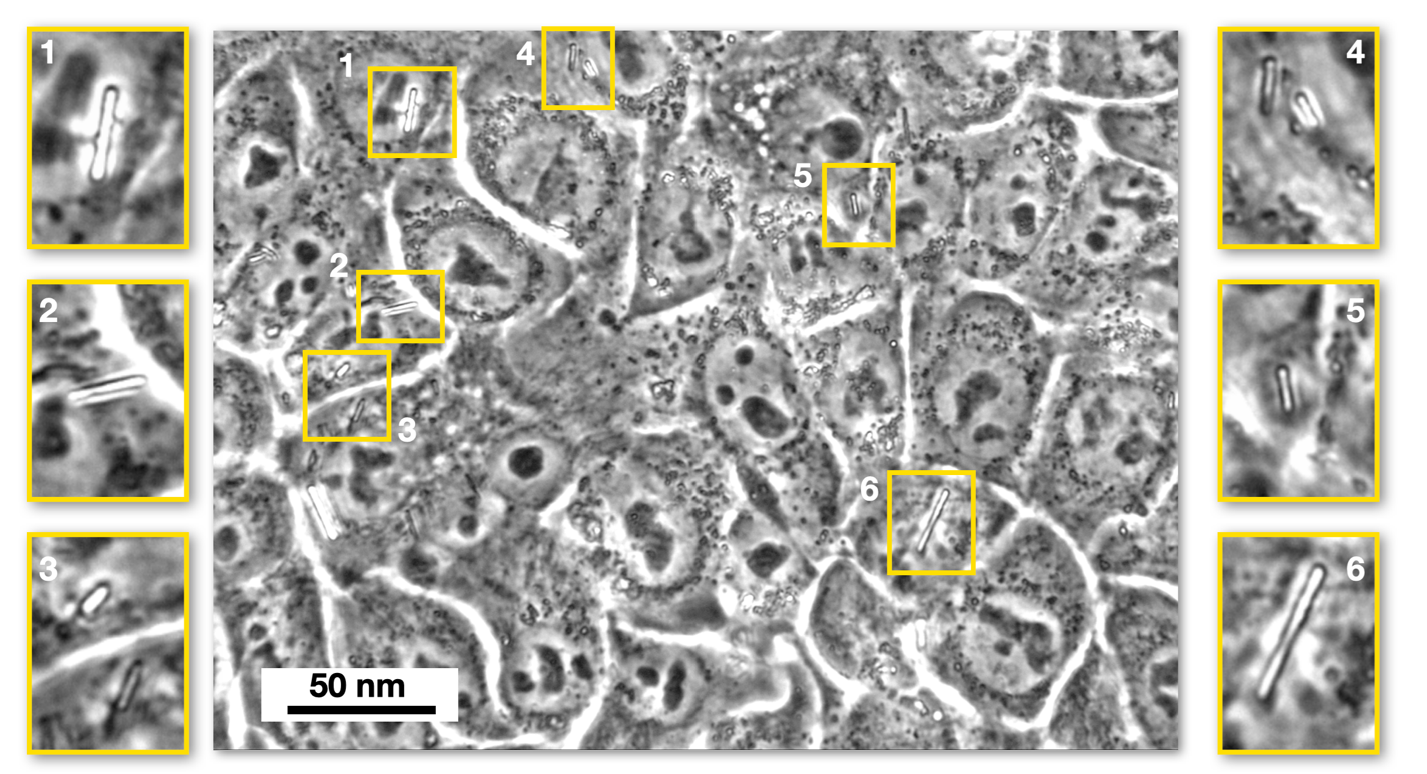

**Figure S2:** The figure presents a microscopy image of a HeLa cell monolayer incubated with magnetic wires, acquired at 60× magnification. Approximately twenty cells are visible, along with fifteen internalized wires. Six fields of view containing magnetic wires, labeled 1 to 6, are shown in enlarged views surrounding the central image. For experiments on living cells, measurements are performed in three distinct fields of view, and the entire procedure is repeated two to three times on separate glass slides prepared under identical culture and incubation conditions. This protocol yields a total of 100 analyzable wires across the full frequency spectrum, allowing for robust determination of the critical frequency $\omega_{C}$ (Eq.1) and the angle $\theta₀$ (Eq. 2) [2].

**Supporting Information S3**

Sampling wires according to their length

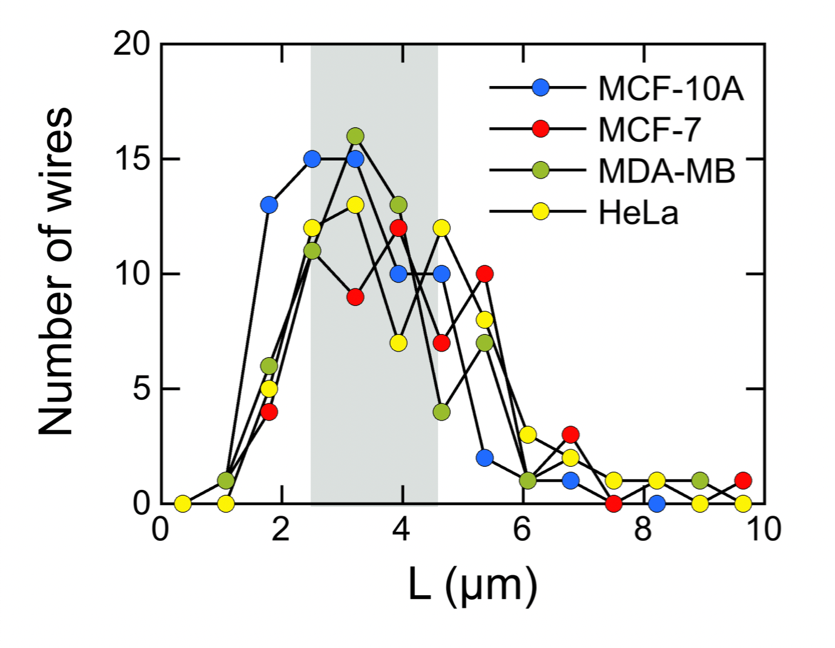

**Figure S3**: Length distribution of wires used in MRS experiments on MCF-10A, MCF-7, and MDA-MB-231 cells. Wires with sizes between 2.5 and 4.5 µm (shaded area) represent 50-70% of the total number.

**Supporting Information S4**

Review of the literature data on the intracellular viscosity obtained for MCF-10A, MCF-7 and MDA-MB-231 breast epithelial cells

| Reference | Microrheology techniques | Passive/Active | Probe size (µm) | Cells | $\boldsymbol{\eta}_{\boldsymbol{0}}$ (Pa s) |
| --- | --- | --- | --- | --- | --- |
| Li & Duits  (2009)[3] | Particle-tracking | Passive | 0.2 | MCF-10A (normal)  MCF-7 (low invasive) | 6.1  0.30 |
| Gal & Weihs  (2014)[4] | Particle-tracking | Passive | 0.2 | MCF-10A (normal)  MCF-7 (low invasive)  MDA-MB-231 (high invasive) | 3.0  2.3  0.41 |
| Guo & Weitz  (2014)[5] | Particle-tracking | Passive | 0.5 | MCF-10A (normal)  MCF-7 (low invasive) | 3.6  0.73 |
| Smelser & Holzwarth  (2015)[6] | Particle-tracking | Passive | 0.4 | MCF-10A (normal)  MCF-7 (low invasive)  MDA-MB-231 (high invasive) | 1.50  1.31  1.26 |
| Mandal & Manneville  (2016) [7] | Optical tweezers | Active | 2.0 | MCF-10A (normal)  MDA-MB-231 (high invasive) | 18  13 |
| Dessard &  Berret (2024)[2] | Magnetic Rotational Spectroscopy | Active | 3.5 | MCF-10A (normal)  MCF-7 (low invasive)  MDA-MB-231 (high invasive) | 42 ± 9  56 ± 17  11 ± 5 |

**Table S1**: Data from the literature on intracellular viscosity in breast epithelial cell lines MCF-10A, MCF-7, and MDA-MB-231. These data can also be found in graphical form in reference [8]. The table also specifies the measurement techniques and probe sizes used. Overall, the data indicate a general trend in which the cancerous MCF-7 and MDA-MB-231 cells exhibit lower intracellular viscosities than the normal-like MCF-10A cells.

**Supporting Information S5**

Comparison between the Newton and Oldroyd-B models based on finite element simulations

The simulations performed with Newton model were carried out using the constitutive equation:

$$\boldsymbol{\sigma}=2\eta_{0}\boldsymbol{D}$$

where $\boldsymbol{\sigma}$ and $\boldsymbol{D}\boldsymbol{=}\frac{\boldsymbol{1}}{\boldsymbol{2}}\left( \boldsymbol{\nabla}\boldsymbol{v}+{\boldsymbol{(\nabla}\boldsymbol{v}\boldsymbol{)}}^{\boldsymbol{T}} \right)$ are the stress and deformation rate tensors, respectively and $v$ the flow velocity.

For the Oldroyd-B model, which describes the behavior of viscoelastic fluids, the constitutive equation is given by [9]:

$$\nabla$$

$$\nabla$$

$$\boldsymbol{\sigma+}\tau\boldsymbol{\sigma}=2\eta_{0}\boldsymbol{(D}{+\tau}_{R}\boldsymbol{D)}$$

$$\nabla$$

where $\boldsymbol{\sigma}$ is the upper-convected time derivative of the stress tensor, $\tau$ and $\tau_{R}$ the relaxation and retardation times. The upper-convected time derivative was introduced to preserve frame invariance in polymer flow and is expressed as:

$$\nabla$$

$$\boldsymbol{\sigma=}\frac{\partial}{\partial t}\boldsymbol{\sigma+}v\cdot\boldsymbol{\nabla}\boldsymbol{\sigma-}\left( {\mathbf{(}\nabla v\boldsymbol{)}}^{\boldsymbol{T}}\boldsymbol{\cdot\sigma+\sigma\cdot}\mathbf{(}\nabla v\boldsymbol{)} \right)$$

In this model, the following relationship $\eta_{0}=\eta_{S}+\eta_{P}$ holds, where $\eta_{S}$ and $\eta_{P}$ are the solvent and polymer viscosities respectively [9]. When $\eta_{S}$ = 0, the constitutive equation of the Oldroyd-B model reduces to the upper-convected Maxwell model.

For the simulations, the Oldroyd-B viscoelastic module integrated into the COMSOL software package was used, with the solvent viscosity $\eta_{S}$ and retardation time $\tau_{R}$ set to zero. The calculations in this section were performed for illustrative purposes on MCF-10A cells, corresponding to a viscosity of $\eta_{0}$ = 46 Pa s and a relaxation time $\tau$ = 0.6 s (Table I). The angular frequency was set to ω = 0.01 rad s⁻¹.

**Figs S5a–c** present, in the upper panels, the two-dimensional projections of the velocity field $(xOy)$, $(yOz)$, and $(xOz)$ planes, respectively. From these figures, the velocity profiles along the $x$*-*, $y$*-*, and $z$*-* directions were extracted and compared with those obtained for a purely viscous fluid governed by Newton constitutive equation. The results show good agreement between the two simulations. It should also be noted that for the viscoelastic model, the mesh size of the simulation domain was increased compared with that used for the viscous fluid. Despite this adjustment, the computation time increased substantially, from approximately 10 minutes for the Newtonian case to about 90 minutes for the Oldroyd-B model, and further rises with increasing frequency.

Given that the flow field remains nearly unchanged except in the immediate vicinity of the simulated wire, partly due to the change in mesh size, we chose to retain the Newtonian model in the main text, as it provides comparable results with a significantly lower computational cost.

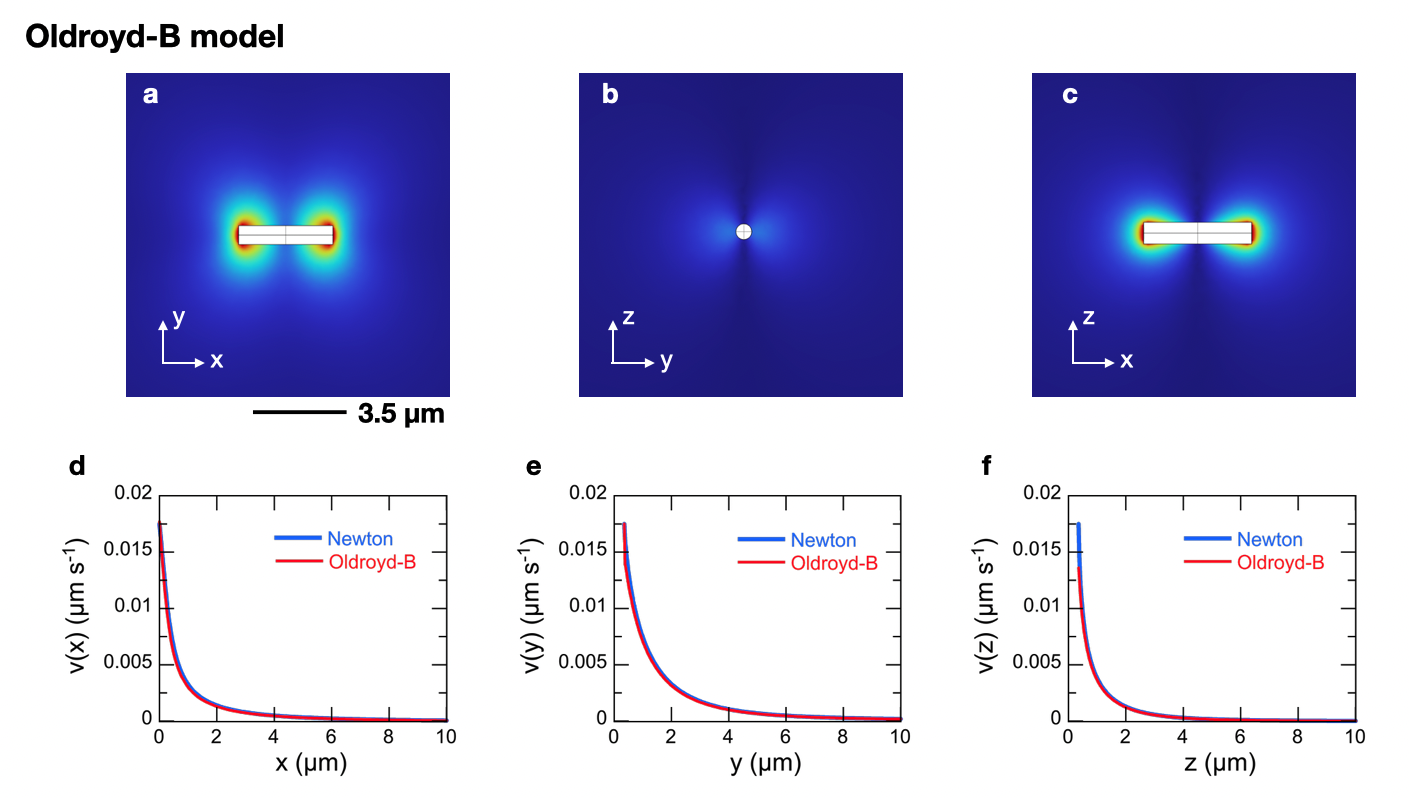

**Figure S5:** **a-c)** 2D-representations of the velocity fields in the $(xOy)$, $(yOz)$, and $(xOz)$ planes, respectively around a rotating wire ($L$ = 3.5 μm, $D$ = 0.70 μm,) obtained from COMSOL finite element modeling using the viscoelastic Oldroyd-B model. The wire rotates at an angular frequency of 0.01 rad s⁻¹ around the $z$-axis, resulting in a maximum tip velocity $v_{Max}$ of 0.0175 μm s⁻¹. **d-f)** Velocity fields as a function of the distance from the wire along the $x$-, $y$-, and $z$-directions respectively for Newton and Oldroyd-B models.

**Supporting Information S6**

Relative variation of the velocity component along $y$- and of the velocity modulus in the $x$-, $y$-, and $z$-directions

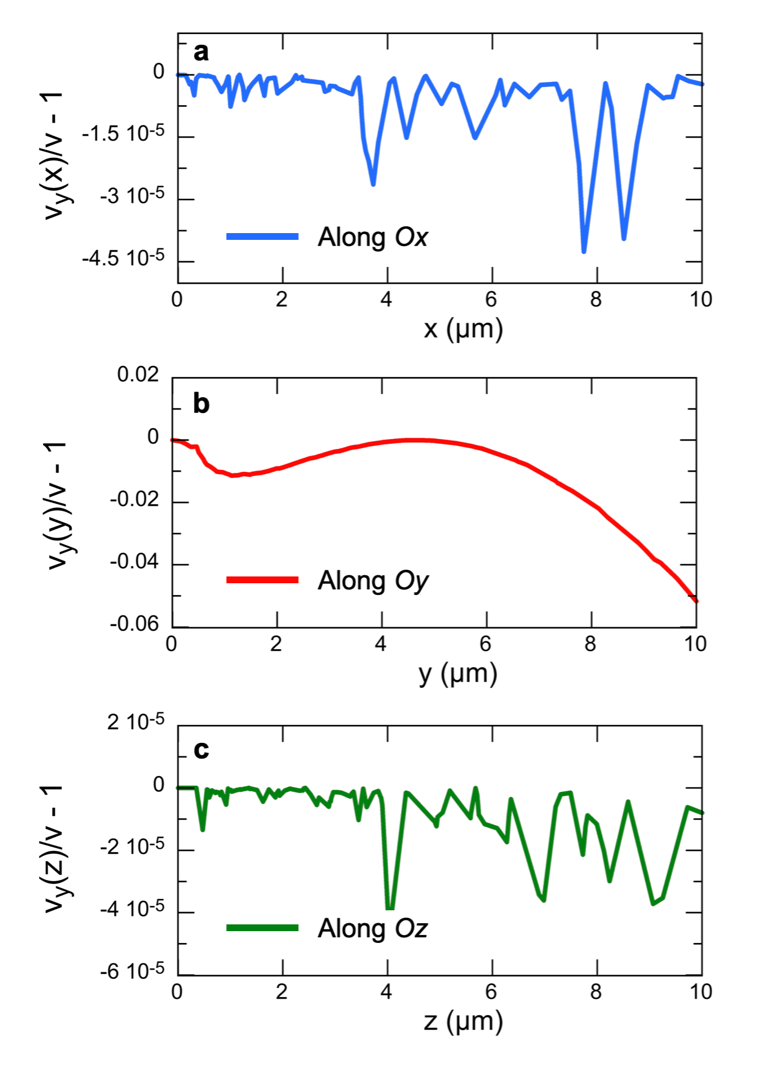

**Figure S6: a)** Relative variation of the velocity component $v_{y}(x)/v(x)-1$ along the $Oy$ direction, and of the velocity modulus $v\left( x \right)=\sqrt{{v_{x}(x)}^{2}+{v_{y}(x)}^{2}+{v_{z}(x)}^{2}}$ as a function of the distance from the wire along the $Ox$ axis. The wire considered in these 3D-finite element COMSOL simulations corresponds to the standard configuration studied in this work, with a length $L$ = 3.5 µm, a diameter $D$ = 0.7 µm, and an angular rotation velocity $\omega_{C}$ = 0.1 rad s⁻¹. b,c) Same as in a) for $v_{y}(y)/v(y)-1$ and $v_{y}(z)/v(z)-1.$

**Supporting Information S7**

COMSOL Multiphysics finite element simulation of rotating wires in the conditions $L/D >>1$

A particularly interesting case is that of an infinitely thin wire, where $L/D >>1$ in rotation in a Newton fluid. As shown below with simulations, this configuration exhibits model behaviors that characterize velocity fields in rotating wires. Notably, universal scaling laws emerge, well described by analytical functions, regardless of wire length. In the 3D simulations provided in the core of the paper, that is with wire diameters closer to $L/D \sim5$, some of these behaviors persist, though they no longer follow simple analytical descriptions.

**Simulation Domain:** Previous studies indicate that meaningful simulations require domains at least 20 times larger than the wire size. For a 20 µm-long wire for instance, simulations span from -200 µm to +200 µm. Here, we conducted simulations for wire lengths of $L$ = 2, 5, 10, 20, and 50 µm with a diameter $D$ = 0.1 µm. Using the COMSOL finite element programs, we used an extremely fine simulation grid to precisely capture the stationary rotational state around the wire.

**Maximum Velocity:** **Figure S7-A** presents the velocity fields in the $(xOy)$ plane for wires of five different lengths, rotating at an angular frequency of $\omega$ = 0.1 rad s^-1^. The axes $Ox$ and $Oy$ are fixed to the coordinate system of the rotating wire. As expected, the maximum velocity $v_{Max}$ occurs at the wire extremity and equal to:

$$v_{Max}=\frac{\omega L}{2}$$

It is important to emphasize that our primary focus is on the velocity components along the $Oy$-axis, as the components along the $Ox$-axis and $Oz$-axis are significantly lower in comparison (**Supporting Information S6**).

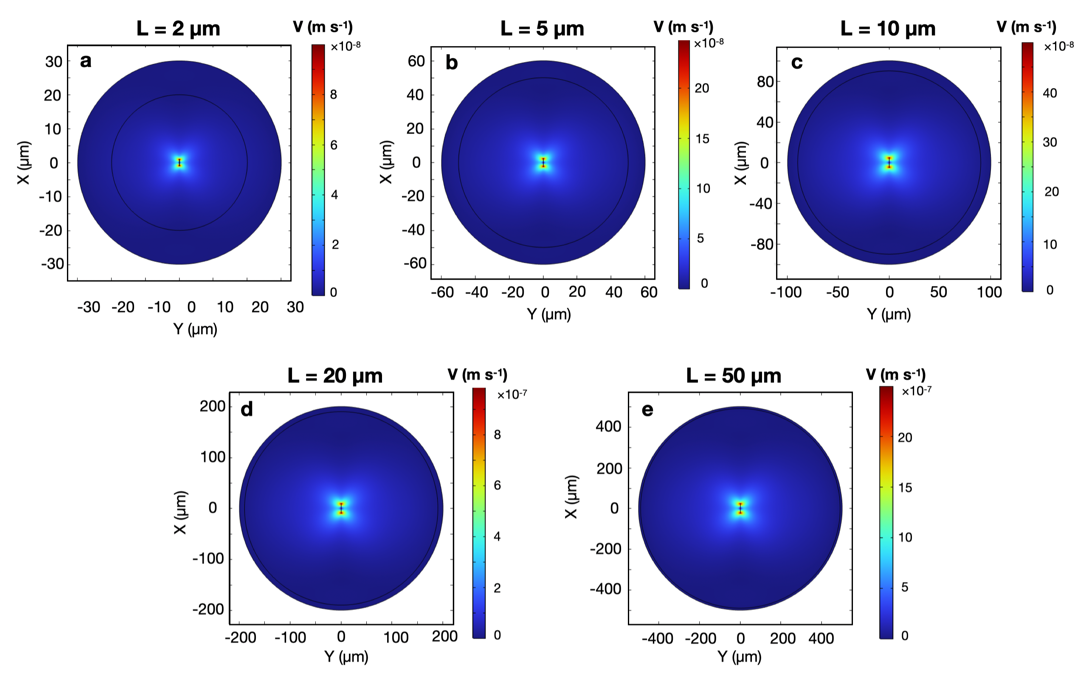

**Figure S7-A:** **a-e)** Velocity fields in the $(xOy)$plane obtained from simulations for wires of lengths $L$ = 2, 5, 10, 20 and 50 µm respectively and diameter $D$ = 0.1 µm.

**Figure S7-B** presents the velocity profiles along the $Ox$- and $Oy$-directions for the five cases studied. In all cases, velocity decreases with distance from the wire. This decay was quantified using a stretched exponential function, revealing two generic scaling laws of the form:

$$v_{y}(x)=\frac{\omega L}{2}exp\left( -\left( x/\delta_{v,x} \right)^{1/2} \right)$$

$$v_{y}(y)=\frac{\omega L}{2}exp\left( -\left( y/\delta_{v,y} \right)^{2/3} \right)$$

where $\delta_{v,x}$ are $\delta_{v,y}$ are the penetration lengths for the velocity in the $Ox$- and $Oy$-directions respectively. The agreement between the simulation data and the stretched exponential fits is satisfactory across the analyzed distance ranges.

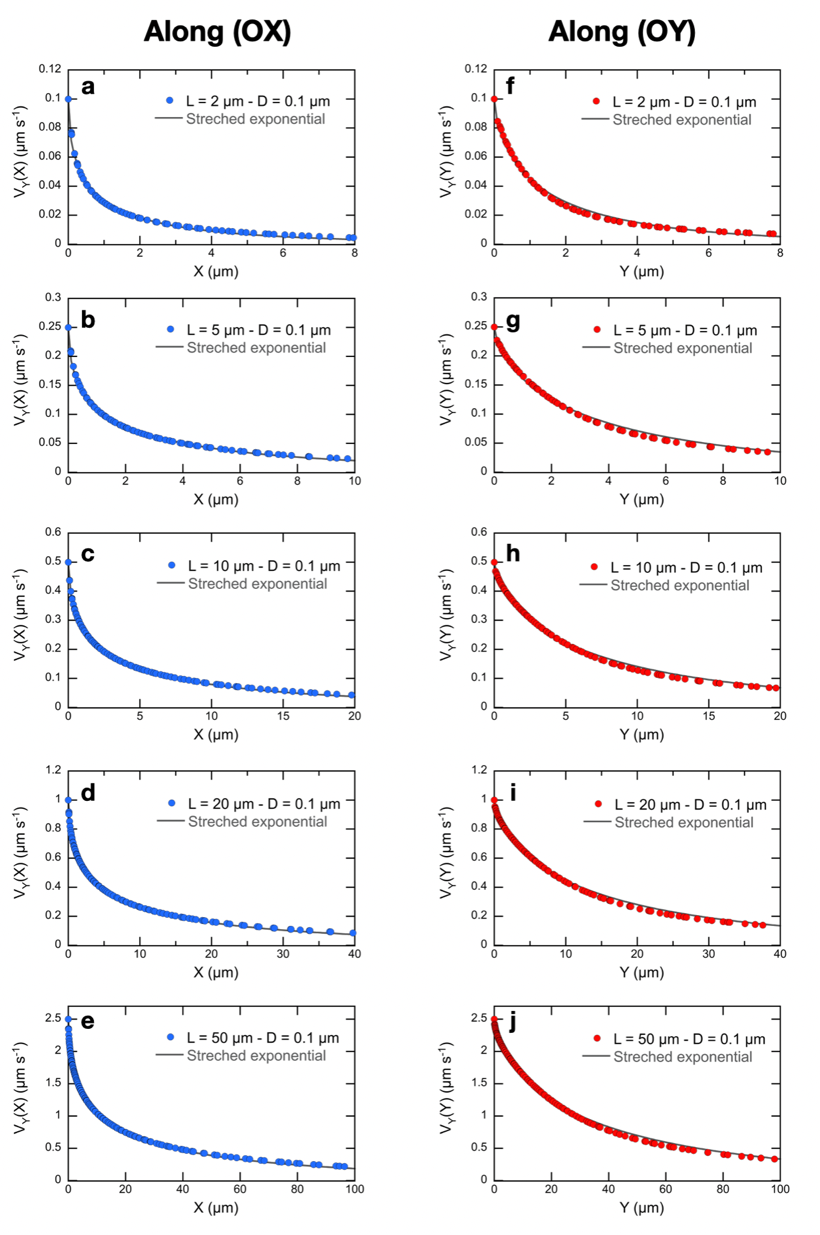

**Figure S7-B: a-e)** Velocity fields along the $Ox$-direction (parallel to the wire) obtained in the conditions of **Fig. S7-A.** The position $x$ = 0 on the diagrams corresponds to the wire end, so for $x = L/2$, $y$ = 0. **f-j)** Same as in a-e) along the $Oy$-direction (perpendicular to the wire). The position *y* = 0 on the diagrams corresponds to the wire end, so for $x = L/2$, $y = D/2$.

**Figure S7-C** presents the variations in penetration lengths $\delta_{v,x}$ and $\delta_{v,y}$ for velocity, as well as $\delta_{\dot{\gamma},x}$ and $\delta_{\dot{\gamma},y}$ for velocity gradients. The results show that penetration lengths scale linearly with $L$, following the relationships:

$$\delta_{v,x}=0.27L;\delta_{v,y}=0.69L$$

These results indicate that the extent of the velocity field increases with wire length and is higher along the $y$-axis. In contrast, the penetration lengths of the velocity gradients remain constant and independent of $L$. For the studied case, we find :

$$\delta_{\dot{\gamma},x}=0.23 \mu m; \delta_{\dot{\gamma},y}=1.23 \mu m$$

Here, it is found that the highest shear zone is localized at the wire ends. In addition, the maximum velocity gradient increases with length, following scaling laws with exponents less than 1:

$$\dot{\gamma}_{Max,x}(L)=0.19L^{0.57}; \dot{\gamma}_{Max,y}(L)=0.057L^{0.43}$$

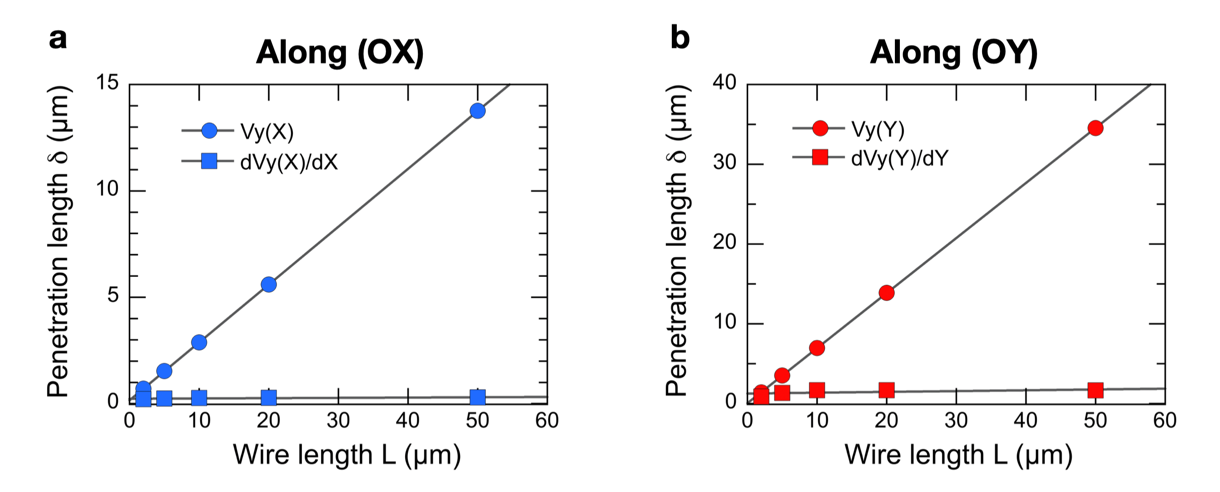

**Figure S7-C**: a) Penetration length $\delta_{v,x}$ and $\delta_{\dot{\gamma},x}$ as a function of the wire length derived from velocity profiles along the $Ox$-direction. b) Same as in a) along the $Oy$*-*direction.

**Supporting Information S8**

Evidence of power-law relationships in MRS data across the studied cell lines for $D(L)$, $L^{*}(L)$, $\omega_{C}(L^{*})$ and $\eta_{0}\left( \omega_{C} \right)$

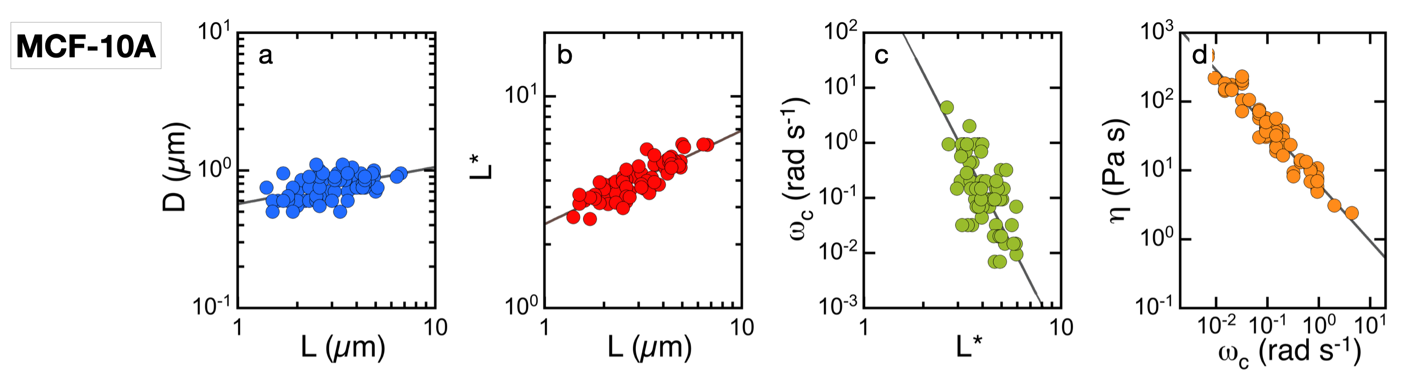

**Figure S8-A:** **a)** Dependence of the diameter of internalized wires in normal-like breast epithelial cells MCF-10A on their length. **b)** Same as in a) for the reduced length $L^{*}=L/\left[ D\sqrt{g(L/D)} \right]$. **c)** Critical frequency $\omega_{C}$ as a function of the reduced wire length $L^{*}$. **d)** Cytoplasmic viscosity $\eta_{0}$ versus the critical frequency $\omega_{C}$. The solid lines in the 4 figures represent least-squares fits to a power law of the form $D(L) \sim L^{\kappa}$, $L^{*}(L) \sim L^{\lambda}$, $\omega_{C}(L^{*}) \sim{L^{*}}^{\mu}$ and $\eta_{0}\left( \omega_{C} \right) \sim{\omega_{C}}^{\nu}$, where $\kappa$, $\lambda$, $\mu$, and $\nu$ are scaling exponents. The coefficients obtained from these fits are listed in **Table S2.**

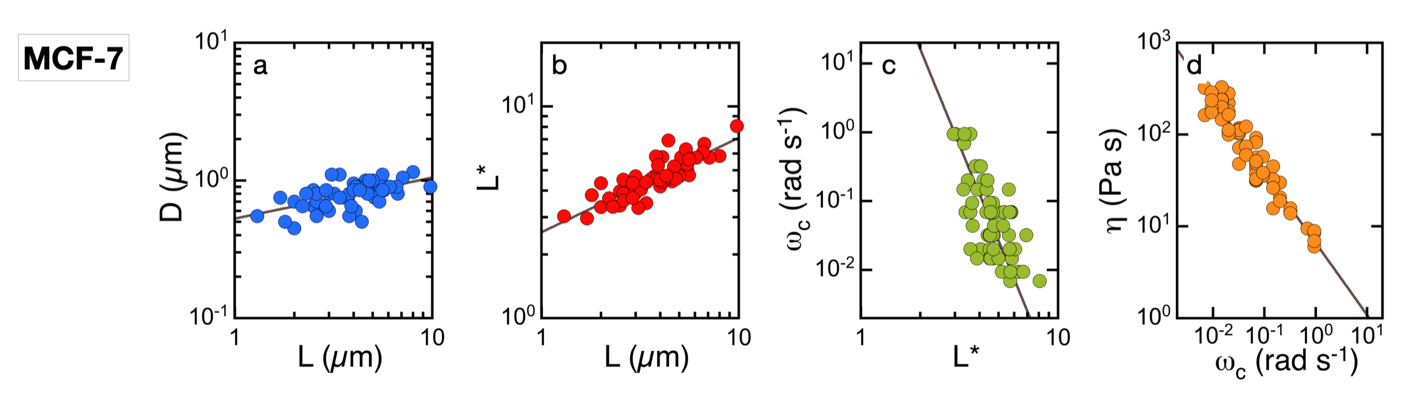

**Figure S8-B:** Same as Fig. S8-A for MCF-7 breast epithelial cells with low a metastatic potential.

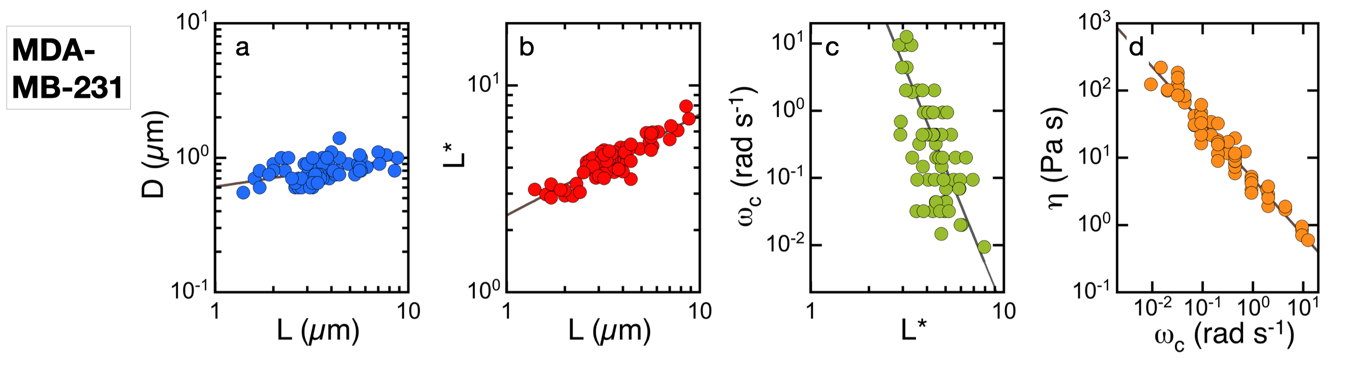

**Figure S8-C:** Same as Fig. S8-A for MDA-MB-231 breast epithelial cells with high a metastatic potential. More details about the data from Figs. S8 A-C can be found in Ref. [2].

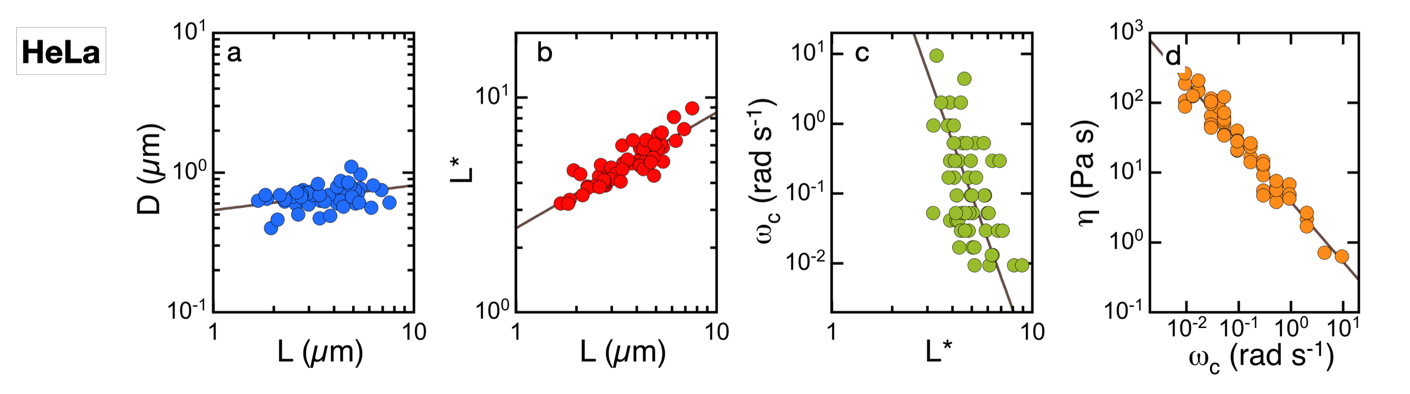

**Figure S8-D**: Same as Fig. S8-A for HeLa cervical cancer cells.

| Cell lines | n | Exponent  $\boldsymbol{\kappa}$ | Exponent  $\boldsymbol{\lambda}$ | Exponent  $\boldsymbol{\mu}$ | Exponent  $\boldsymbol{\nu}$ |
| --- | --- | --- | --- | --- | --- |
| MCF-10 | 68 | 0.267 | 0.442 | -7 | -0.827 |
| MCF-7 | 60 | 0.298 | 0.446 | -7 | -0.781 |
| MDA-MB-231 | 68 | 0.218 | 0.488 | -7 | -0.833 |
| HeLa | 55 | 0.177 | 0.539 | -8 | -0.859 |

**Table S2:** Summary of the number of wires analyzed and the power-law exponents obtained for each of the four cell types shown in Figs. S8 A–D.

References

[1] M. Radiom, E.K. Oikonomou, A. Grados, M. Receveur, J.-F. Berret, Probing DNA-Amyloid Interaction and Gel Formation by Active Magnetic Wire Microrheology, in: V. Arluison, F. Wien, A. Marcoleta (Eds.) Bacterial Amyloids: Methods and Protocols, Springer US, New York, NY, 2022, pp. 285-303.

[2] M. Dessard, J.-B. Manneville, J.-F. Berret, Cytoplasmic viscosity is a potential biomarker for metastatic breast cancer cells, *Nanoscale Adv.* **6**, 1727-1738 (2024). doi: 10.1039/d4na00003j

[3] Y. Li, J. Schnekenburger, M.H. Duits, Intracellular Particle Tracking as a Tool for Tumor Cell Characterization, *J. Biomed. Opt.* **14**, 064005 (2009). doi: 10.1117/1.3257253

[4] N. Gal, D. Weihs, Intracellular Mechanics and Activity of Breast Cancer Cells Correlate with Metastatic Potential, *Cell. Biochem. Biophys.* **63**, 199-209 (2012). doi: 10.1007/s12013-012-9356-z

[5] M. Guo, A.J. Ehrlicher, M.H. Jensen, M. Renz, J.R. Moore, R.D. Goldman, J. Lippincott-Schwartz, F.C. Mackintosh, D.A. Weitz, Probing the Stochastic, Motor-Driven Properties of the Cytoplasm Using Force Spectrum Microscopy, *Cell* **158**, 822-832 (2014). doi: 10.1016/j.cell.2014.06.051

[6] A.M. Smelser, J.C. Macosko, A.P. O'Dell, S. Smyre, K. Bonin, G. Holzwarth, Mechanical Properties of Normal versus Cancerous Breast Cells, *Biomech. Model. Mechanobiol.* **14**, 1335-1347 (2015). doi: 10.1007/s10237-015-0677-x

[7] K. Mandal, A. Asnacios, B. Goud, J.-B. Manneville, Mapping Intracellular Mechanics on Micropatterned Substrates, *Proc. Natl. Acad. Sci.* **113**, E7159-E7168 (2016). doi: 10.1073/pnas.1605112113

[8] A.M. Markl, D. Nieder, D.I. Sandoval-Bojorquez, A. Taubenberger, J.F. Berret, A. Yakimovich, E.S. Oliveros-Mata, L. Baraban, A. Dubrovska, Heterogeneity of tumor biophysical properties and their potential role as prognostic markers, *Cancer Heterogen. Plasticity* (2024). doi: 10.47248/chp2401020011

[9] R.G. Larson, *Constitutive Equations for Polymer Melts and Solutions*, Butterworth-Heinemann1988.
